# Spatial specificity of MADS-box transcription factors in floral organs

**DOI:** 10.64898/2025.12.18.695104

**Authors:** Emma Désert, Michiel Vandenbussche, Marie Monniaux

## Abstract

**Highlight:** MADS-box transcription factors specify the identity of floral organs at the primordia stage, but they also have late and spatially-restricted roles in developing or mature floral organs, which we explore in this review.

Floral homeotic genes, most of them encoding MADS-box transcription factors, are classically pictured as early specifiers of floral organ identity, as implied by the famous ABC model. Yet, floral homeotic genes remain expressed throughout floral organ development, sometimes with a cell type-specific expression pattern, and late functions have been firmly established for some of these genes. Here, we review how floral homeotic MADS-box genes are expressed in floral organs during their development, highlighting general trends for their spatial or temporal-specific expression, in particular for B-class genes in petals that are systematically distally-enriched. We focus on chosen examples from the literature to discuss the different roles associated with this specific expression, and we then explore the possible molecular mechanisms by which MADS-box transcription factors can adopt spatially or temporally restricted functions in floral organs, either by a simple restriction of their presence, or by a different mode of action in given cell types. Altogether, this review highlights that late roles of floral MADS-box transcription factors have been largely unexplored, and that these regulators might have multiple different functions according to the cell types in which they are present.

## Introduction

Ask the readers of the Flowering Newsletter about MADS-box genes, and you would likely hear about their crucial role in plant development and reproduction. Ask them about their expression pattern, and surely the ABC model of floral organ identity would be mentioned, in which the identity of floral organs is defined by the whorl-specific and combinatorial expression of (mostly) MADS-box genes. Ask them again what MADS-box genes later do in a developing floral organ, across cells and tissues, and until the organ is mature, and the answer might not be so clear. Indeed, our limited knowledge of the late functions of homeotic transcription factors reflects a fundamental limitation of classical loss-of-function mutant analyses, which perturb early developmental processes and thus obscure their roles at later stages. In contrast to their well-described role in the early specification of floral organ identity, the late role of MADS-box genes, accompanied by their temporally- and spatially-defined expression during floral organ development, is much less characterized and is the purpose of this review.

The ABC model is a genetic framework for floral organ identity specification, relying on the combinatorial expression of genes belonging to A-, B- and C-classes: A-class genes expression alone specifies sepal identity, A+B-class genes expression specifies petal identity, B+C-class genes expression specifies stamen identity, and C-class genes expression specifies carpel identity (Schwarz-Sommer *et al*., 1990; Coen and Meyerowitz, 1991; Bowman and Moyroud, 2024). While this model has been refined over the years, by the addition of the ovule identity and floral identity D- and E-classes respectively (Colombo *et al*., 1995; Pelaz *et al*., 2000; Ditta *et al*., 2004), the debate on the existence of the A-function and its molecular conservation (Cartolano *et al*., 2007; Causier *et al*., 2010; Morel *et al*., 2017, 2019), and the multiple variations on the strict ABC-model with fading borders, sliding borders, or all kinds of sub- or neo-functionnalization events (Soltis *et al*., 2007; Bowman and Moyroud, 2024), this model still holds to define floral organ identity in Angiosperms, across their stunning morphological diversity. Originally, the ABC model was supported by two types of observations in *Arabidopsis thaliana* (Arabidopsis) and *Antirrhinum majus* (Antirrhinum): mostly by phenotypes of floral homeotic mutants, but also by their restricted expression patterns in the floral meristem, resulting in the whorl-specific definition of floral organs (Schwarz-Sommer *et al*., 1990; Coen and Meyerowitz, 1991). Almost all of the ABC-class genes belong to the large family of MADS-box transcription factors, in particular from the type II MIKC-C class (Gramzow and Theissen, 2010). Their specific expression pattern is the result of transcriptional regulation by a combination of spatially-localized transcription factors. For instance in Arabidopsis, LEAFY (LFY) and FLOWERING LOCUS T (FT) independently activate the A-class gene *APETALA1* (*AP1*) throughout the floral meristem (Parcy *et al*., 1998; Wagner *et al*., 1999), but its expression will be repressed in the center of the floral meristem by the C-class gene *AGAMOUS* (*AG)* (Gustafson-Brown *et al*., 1994), whose expression is itself activated in this domain by LFY, WUSCHEL (WUS) and SEPALLATA3 (SEP3) (Lohmann *et al*., 2001; Liu *et al*., 2009). The B-class genes *APETALA3* (*AP3*) and *PISTILLATA* (*PI*) are expressed in the petal and stamen whorls by the combined action of LFY and UNUSUAL FLORAL ORGANS (UFO), the later being only expressed in this domain (Lee *et al*., 1997; Rieu *et al*., 2023). Additionally, boundary genes like *SUPERMAN* refine the precise expression domains of ABC-class genes and thereby the correct location and size of organ whorls (Sakai *et al*., 1995; Prunet *et al*., 2017). Once their spatially localized transcription is established, it is generally assumed that translational regulation is limited, and that MADS-box transcription factors only move to a limited range, if at all, depending on the protein (Perbal *et al*., 1996; Urbanus *et al*., 2009, 2010); however this has not been examined systematically and it will be discussed later in this review. MADS-box proteins are also known to form tetrameric complexes, as proposed in the quartet model, which is the molecular support for their combinatorial mode of action (Honma and Goto, 2001; Theißen and Saedler, 2001; Käppel *et al*., 2023). They also interact with protein partners from other families of regulators, in particular AUXINE-RESPONSE FACTORS (ARFs), SQUAMOSA Promoter-Binding Protein-Like proteins (SPLs), or KNOTTED1 (KN1)-like homeobox proteins (KNOX) (Smaczniak *et al*., 2012; Bemer *et al*., 2017; Abraham-Juárez *et al*., 2020; Goslin *et al*., 2023; van Mourik *et al*., 2023), and the exact function of these molecular interactions has been established in a small number of cases (Smaczniak *et al*., 2012; José Ripoll *et al*., 2015; Bemer *et al*., 2017).

Homeotic genes like MADS-box genes might have been initially viewed as the first specifiers of organ identity, merely acting at an initial stage through an early pulse of expression, with downstream players taking over to build fully developed organs. However, it is now well known that long after floral organ identity has been defined, floral homeotic MADS-box transcription factors continue to be expressed. For instance, *AG* is still expressed in stamens and carpels at late stages, but in specific cell types only (tapetum and endothecium in stamens, stigmatic papillae and ovular endothecium in the carpel, and nectaries) (Bowman *et al*., 1991; Ito *et al*., 2007). A late expression (*i.e.* after the floral organ primordia stage) has been reported for many floral MADS-box transcription factors across several studies, sometimes in association with a specific role, but we lack an integrated view of this process. In this review, we attempted to compile the clearest examples of spatially- and temporally-restricted expression of floral MADS-box genes throughout floral organ development, and we describe a small number of cases when a functional role for this specific expression has been established. We then discuss the possible molecular mechanisms that could lead to the spatialization of function of floral MADS-box genes.

### Spatially restricted expression of MADS-box genes within floral organs

In the following paragraphs, we compiled examples of detailed expression patterns for floral MADS-box genes in developing floral organs, either at the transcript or protein level, stemming from different experimental assays.

#### Localizing MADS-box transcripts

Several techniques allow to explore the expression pattern of floral MADS-box genes, either by locating the transcript or the protein, with different levels of spatial and quantitative precision. The most straightforward assay is real-time quantitative PCR (RT-qPCR), with a limited level of spatial resolution that can be achieved by dissecting the floral organs. This was for instance the approach chosen by Hsu *et al*., (2021) in orchid flowers (*Orchidaceae*), that display a complex perianth composed of sepals, petals and a modified petal with a particular shape, called the lip or labellum. In *Oncidium* orchid flowers, RT-qPCR on lip, petal and sepal organs, revealed that B- and E-class MADS-box genes each had a different expression pattern, specific to some perianth organs (Figure 1A, Hsu *et al*., 2015). This observation, together with functional assays, led to the establishment of the perianth-code model, in which a balance of E-class (OAGL6-1/2) and B-class (OPI and OAP3-1/2) MADS proteins form different complexes determining lip, petal or sepal identity: the OAP3-1 + OAGL6-1 complex specifies the petal or sepal identity (SP complex) while the OAP3-2 + OAGL6-2 complex specifies the lip identity (L complex), both complexes also comprising the other B-class protein OPI. This model was later refined by the identification of additional complexes in *Phalaenopsis* orchids (Hsu *et al*., 2021): the SP’ complex (OAGL6-1+OAP3-2) that is expressed only in petals, and the L’ complex (OAGL6-2 + OAP3-1) that is expressed only in the lower part of lateral sepals. Here, the level of precision achieved by macroscopic dissection of floral organs and RT-qPCR was sufficient to uncover a spatial specificity of MADS-box transcripts within single floral organs; however this approach remains limited to flowers with large organs and at late stages of development, accessible to dissection.

**Figure 1.**
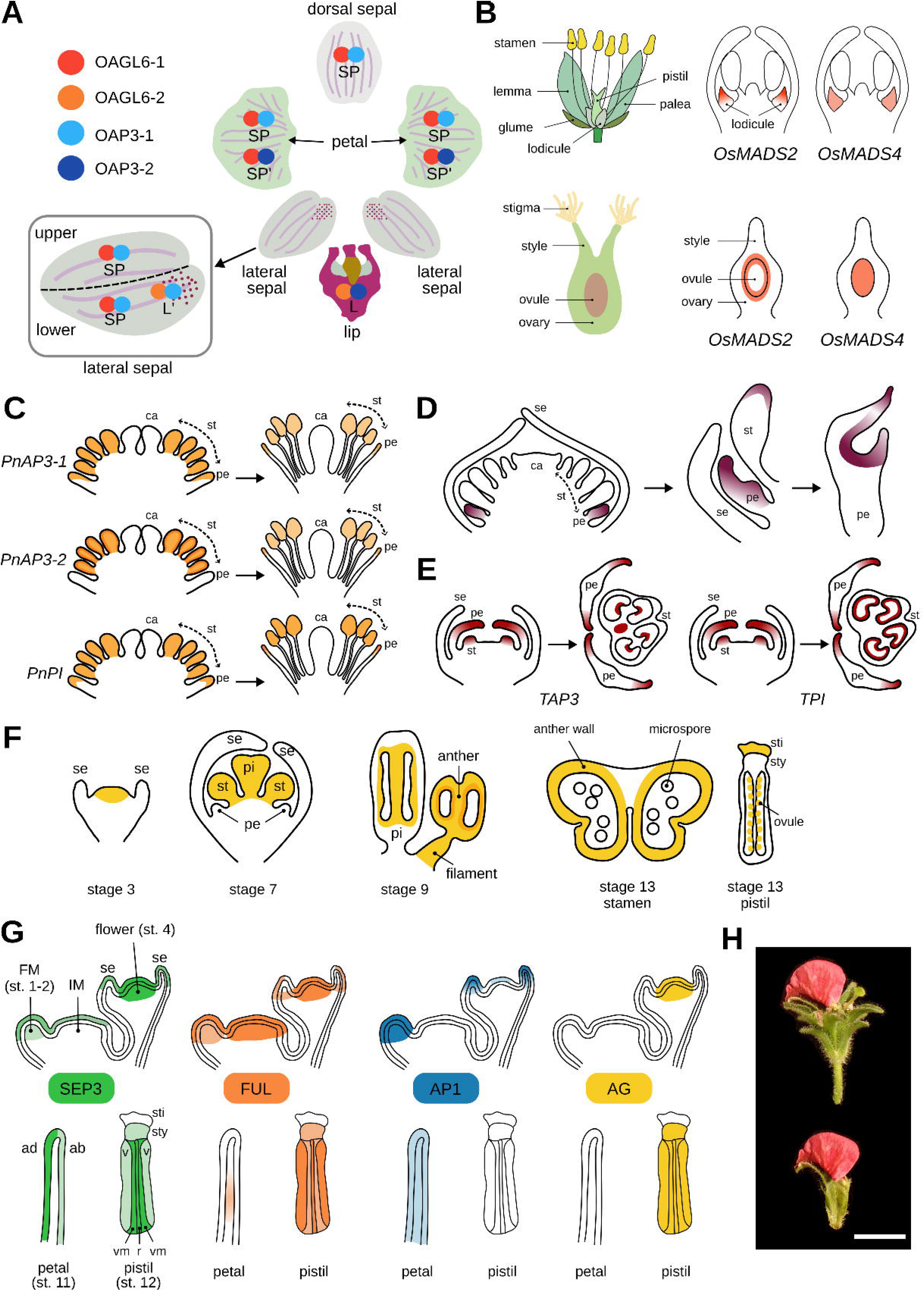
Chosen examples of floral MADS-box transcripts or proteins with a precise spatial localization described in developing or mature floral organs. **(A)** Orchid *Phalaenopsis* perianth-code. The dorsal sepal, petal, lateral sepal and lip express different combinations of OAGL6-1, OAGL6-2, OAP3-1 and OPA3-2 proteins, forming complexes with OPI (not displayed here). The lateral sepal displays a further zonation into its upper and lower part, expressing different protein complexes. Adapted from Hsu *et al*., 2021. **(B)** During rice floret development, *OsMADS2* is distally and laterally enriched in the lodicules, which restricts lodicule elongation (for simplicity, only expression in the lodicule is depicted here). In the pistil, *OsMADS2* expression is enriched in the ovary adaxial layers and ovule outermost layers. *OsMADS4*, in contrast, is uniformly expressed in the lodicule and in the ovule. The expression patterns are summarized from Yadav *et al*., 2007; Zamzam *et al*., 2025, and for simplicity, only the expression in the organs of interest (lodicules and pistil) are depicted here. Floret and pistil schemes are redrawn from Shen *et al*., 2021. **(C)** Simplified expression patterns of *PnAP3-1*, *PnAP3-2* and *PnPI* in flowers of *Papaver nudicaule* at two developmental stages (adapted from Kramer and Irish, 1999). For simplicity, only the expression in the organs of interest (petals and stamens) is depicted here. **(D)** Simplified expression pattern of *NdAP3-3* in flowers of *Nigella damascena* at three developmental stages (adapted from Zhang *et al*., 2013). **(E)** Simplified expression pattern of *TAP3* and *TPI* in flowers of *Solanum lycopersicum* (tomato) at two developmental stages and in a longitudinal (left) or cross (right) section. Adapted from Martino *et al*., 2006. **(F)** Simplified expression pattern of *AG* during *Arabidopsis thaliana* flower development, based on *in situ* hybridization results from (Bowman *et al*., 1991a; Ito *et al*., 2004). **(G)** Simplified patterns of translational fluorescent reporters for SEP3, FUL, AP1 and AG during *Arabidopsis thaliana* flower development, showing the inflorescence meristem (IM) with a flower meristem (FM) at stage 1-2 and a stage-4 flower initating its sepals, a stage-11 petal and a stage-12 pistil. Adapted from (Urbanus *et al*., 2009. **(H)** *Petunia hybrida phdef-151* flower with a revertant sector restoring PhDEF function in the epidermal layer (top: full flower; bottom: isolated sepal). Late epidermal expression of *PhDEF* is sufficient to restore limb shape and pigmentation. Scale bar = 1 cm. se = sepal, pe = petal, st = stamens, ca = carpel, pi = pistil, sti = stigma, sty = style, ad = adaxial side, ab = abaxial side, v = valve, vm = valve margin, r = replum. For all expression patterns reported here, the color shades reflect the strength of expression.

*In situ* hybridisation (ISH) is commonly used to assess spatial localization of transcripts on floral organ sections; hence possibly at very young stages of development and with a high spatial resolution. In rice (*Poaceae*), ISH revealed that the two *PI/GLO* orthologs *OsMADS2* and *OsMADS4* have a different expression pattern in the lodicules, the second-whorl organs of the rice spikelet: *OsMADS4* is expressed in a uniform manner, while *OsMADS2* transcripts accumulate in the distal and peripheral lodicule region during late floret development (Figure 1B, Yadav *et al*., 2007). Similarly, while *OsMADS4* is evenly expressed in the ovary and ovules, *OsMADS2* expression is restricted to the adaxial layers of the ovary and the outermost layers of the ovules (Figure 1B, Zamzam *et al*., 2025). In *Papaver nudicaule (*Iceland poppy, *Papaveraceae)*, *PnAP3-1* and *PnPI-1* transcripts are detected by ISH homogeneously in the petal primordium, then they are gradually restricted to the distal part of the petals when stamen primordia start to emerge, while *PnAP3-2* transcripts are only detected at late stages of petal development and restricted to the tip of the petal (Kramer and Irish, 1999, Figure 1C). In stamens, *PnAP3-1/2* and *PnPI-1* genes are uniformly expressed, with an epidermal enrichment for *PnAP3-2.* Similarly, *Dicentra eximia* (fringed bleeding-heart, *Papaveraceae*) *DePI* and *DeAP3* transcripts are uniformly expressed throughout stamen development while their expression is not detected in petal primordia, and only detected for *DePI* in the epidermis and vasculature of the petal base during later flower development (Kramer and Irish, 1999). In the highly derived petals of *Nigella damascena* (*Ranunculaceae*), the B-class gene *NdPI* displays an even expression in the petal primordia, while *NdAP3-2* and *NdAP3-3* are more strongly expressed with a distal enrichment (Figure 1D). *NdAP3-3* expression persists when the petal develops and its transcript accumulates at the crease of the petal, where the nectary chamber will form (Gonçalves *et al*., 2013; Zhang *et al*., 2013; Yao *et al*., 2019). Spatially restricted expression of *AP3/PI* homologs is also observed in tomato (*Solanaceae*): at early stages of flower development *TAP3*, *TM6* and *TPI* expression is uniform in petal and stamen primordia, but it becomes gradually restricted to the lateral edges and distal part of petals for *TAP3* and *TPI*, as well as vascular bundles and tapetal cells of stamens for *TAP3* and *TPI*, and ovule inner integument for *TM6* (Figure 1E, Martino *et al*., 2006).

A dynamical expression pattern has been described for the C-class gene *AG* in Arabidopsis by ISH (Bowman *et al*., 1991; Ito *et al*., 2004) (Figure 1F). Initially, *i.e.* at stages 1-6 for stamens and stages 1-8 for carpels (Smyth *et al*., 1990), *AG* mRNA signal is uniform, but it becomes later restricted to specific cell types. At early stage 9, *AG* mRNA is detected in filament cells and anther walls, but at stage 13, *AG* mRNA signal is left only in the anther walls (endothecium and decaying epidermis), and always absent from the microspore lineage (future pollen grains). In the carpel, *AG* mRNA signal is high in the ovule primordia but low in ovary wall tissues at stage 9, but fades in the style, ovary and placenta when completely formed at stage 12, while remaining high in ovules and stigmatic tissue. In fully developed carpels at stage 14, *AG* mRNA is restricted to the endothelium and stigmatic papillae. *AG* expression is also detected in nectaries, located at the base of the pistil, throughout flower development (Bowman *et al*., 1991; Morel *et al*., 2018). In summary, *AG* displays a complex and dynamical expression pattern during stamen and pistil development, with a restricted expression in specific cell types when floral organs are mature. It is to be noted that this late expression pattern is independent from the repressive action of the A-class gene *AP2*, since the *ap2-1* weak mutant allele also displays *AG* expression in stigmatic papillae and rudimentary ovules (Bowman *et al*., 1991), while the early expression of *AG* in the carpel primordia is indeed dependent on *AP2* repressive action. To our knowledge, it is unknown how this restricted expression is controlled, but this pattern is linked to a specific role of *AG* in regulating microspore (non-cell-autonomously), nectary and ovule development (Ito *et al*., 2004; Morel *et al*., 2018).

Although ISH is a method of choice for the precise description of gene transcriptional patterns in any given species, it can remain technically challenging to set up, and does not work reliably on floral organs at a very late stage of development, in which cells are big and transcript amounts are low. Spatial transcriptomics, including techniques such as single-cell or -nuclei RNA-Seq or Slide-Seq (genome-wide detection of gene expression on tissue sections), are extremely promising avenues to finely characterize floral transcripts in individual cell types (Nolan and Shahan, 2023). They are in theory accessible to any non-model organism with a sequenced genome, although these assays are expensive and the technical and analytical aspects remain challenging.

#### Localizing MADS-box proteins

The previously described methods are used to follow the expression patterns of MADS-box genes, but cannot assess protein localisation following translation and potential protein movement. Immunohistochemistry (IHC, also called immunolocalization) is an assay that relies on specific antibodies directed against the protein of interest, to detect its presence on tissue sections directly. This technique is amenable to species recalcitrant to transformation, but relies on the production of specific antibodies (which is not necessarily straightforward for floral MADS-box proteins with a highly conserved MADS domain), is labour-intensive and risky; therefore only a small number of studies have reported the use of this technique to detect MADS-box proteins localization in floral organs. In the previously cited *Papaveraceae* species *Papaver nudicaule* and *Dicentra eximia*, IHC was performed with polyclonal antibodies raised against AP3 or PI proteins from Arabidopsis (Kramer and Irish, 1999). This showed that overall, the AP3 or PI proteins localization followed the sum of their mRNA patterns in these species. Indeed, PnAP3-1/2 and PnPI-1 were clearly detected in stamens and restricted to the edge of the petals, while DePI and DeAP3 were found homogeneously in stamens, in the placental region of carpels and at the base of the petal (vasculature), as well as in the petal epidermis for DePI only. In these cases, the protein expression pattern was consistent with the gene expression pattern, indicating that there is no significant post-translational regulation or protein movement (Kramer and Irish, 1999).

For species amenable to genetic transformation, one can follow endogenous protein localisation with translational reporters, *i.e.* by fusing a fluorescent protein to the protein of interest. The choice of location of the fusion is crucial since it might disrupt protein function (for DNA binding or interaction with protein partners), therefore making it crucial to validate the protein fusion functionality by complementation in a mutant background. For example, the AP1-GFP (C-term fusion) can rescue the *ap1-15* phenotype while the GFP-AP1 (N-term fusion) construct cannot, likely because the GFP is too close to the DNA binding domain of AP1 located in N-term (Wu *et al*., 2003). Using the full genomic region of the MADS-box gene of interest is also key to avoid co-suppression, a frequent outcome when using ubiquitous promoters like *35S*, and to ensure a specific expression pattern (Urbanus, 2010). Urbanus *et al*. (2009) investigated the *in planta* localisation patterns of four MADS-box proteins in Arabidopsis: AP1 (A-class), AG (C-class), SEP3 (E-class) and FRUITFULL (FUL, specifying floral meristem identity and playing a role in carpel and fruit development (Ferrándiz *et al*., 2000a,*b*) by constructing functional C-term GFP fusion lines. This revealed an expression profile more complex and dynamic than anticipated from their mRNA expression patterns (summarized in Figure 1H). In the inflorescence meristem, SEP3 was only detected in the epidermis while FUL was found in all cell layers. In the flower meristem (stages 1-2, Smyth *et al*., 1990), epidermal SEP3 expression was maintained with occasional expression in inner layer cells, while FUL expression decreased (mainly in inner layers) due to the negative regulation by AP1, whose expression was visible in all cell layers. At stages 4-5, SEP3 expression was maintained in the sepal abaxial and adaxial epidermis, as well as in the center of the floral meristem. AP1 was expressed in sepal tips and petal primordia, AG was expressed at the center of the meristem, and FUL was expressed in the basal region of sepals in all cell layers, but only in the epidermis in their apical part. In the petals from stage 5 to 12, SEP3 retained its epidermal expression and from stage 9 onwards, it was strikingly enriched in the adaxial epidermis over the abaxial epidermis. AP1 expression in all cell layers of the petal was progressively reduced, and FUL expression was only detected at the center of the petal claw. In the ovary wall at stage 12, SEP3 and FUL were both expressed in the replum, valve margins and valve, but in different cell layers, while AG was expressed in all layers of the ovary wall. These complex expression patterns, partially summarized in Figure 1H, suggest a finely tuned regulation of transcription, translation and/or protein movement, which might support the cell type- and cell layer-specific roles of these MADS-box proteins, for the moment mostly uncharacterized.

#### General patterns of MADS-box preferential expression?

From this dispersed evidence of MADS-box transcript or protein expression patterns, in different organs, tissues, species and developmental stages, can we draw general principles? Our observations highlight that, broadly speaking, MADS-box transcripts and/or proteins are often enriched in the distal part of developing organs at advanced stages of development (which often corresponds to the region of the organ that is under differentiation). This is particularly striking for B-class genes/proteins in petals, for which distal (and to some extent epidermal) enrichment seems to be the rule. Indeed, as previously described, it is the case in petals of *Papaver nudicaule* and *Dicentra eximia (Papaveraceae),* but also in *Ranunculus ficaria* (*Ranunculaceae*), tepals of *Michelia figo* (*Magnoliaceae*) and *Sagittaria Montevidensis* (*Alismataceae*, Monocots), and petals of Arabidopsis (*Brassicaceae*), Antirrhinum (*Plantaginaceae*), petunia and tomato (*Solanaceae*) (Perbal *et al*., 1996; Kramer and Irish, 2000; Jenik and Irish, 2001; Martino *et al*., 2006; Chopy *et al*., 2023), among others. The broad phylogenetic distribution of this pattern suggests it is likely ancestral. In Arabidopsis, SEP3 also displays distal and epidermal enrichment in the petals, with a stronger expression in the adaxial epidermis (Urbanus *et al*., 2009), but whether this pattern is general for SEP-like proteins remains to be tested. If confirmed, it would be tempting to speculate that this particular expression pattern of B-and E-class proteins could trigger the development of the very specific petal epidermal features such as conical cells, pigmentation or volatile emission, which are often a hallmark of the adaxial petal epidermis, and typically developing at the tip of the petal rather than at its base. In stamens and pistils, we have not identified a preferential expression pattern of MADS-box genes in developing organs, from the studies that we surveyed. However, we acknowledge that our focus has been more intense on B-class MADS-box genes expression in petals, and that a preferential expression pattern for A-, C-, D-and E-class genes in sepals, stamens and carpels would deserve further attention.

### From localized expression to localized function

Although evidence for uneven expression of floral MADS-box genes in developing floral organs has been accumulating, the exact function associated with this localized expression remains unknown in most cases. Indeed, since knock-outs or knock-downs generally perturb floral organs from the stage of their initiation, late roles of MADS-box genes in restricted spatial domains cannot always be confidently evaluated with these approaches. This is the case of the previously described petals of *Nigella damascena*, where *NdAP3-3* expression is specific to the petal and enriched first distally, then at the inner crease of the petal. An apetalous morph is associated with the loss of *NdAP3-3* expression (likely caused by a transposable element insertion in the second intron of the gene), and knocking-down *NdAP3-3* expression by VIGS causes homeotic transformation of the petals into sepals (Gonçalves *et al*., 2013; Zhang *et al*., 2013). This shows that *NdAP3-3* is necessary for the correct specification of petal identity, but its particular role in the inner crease of the petal, that might be in link with nectary chamber formation, is not accessible through an early knock-down approach. On the contrary, if organ identity is specified redundantly by several MADS-box genes, knocking-out or - down a single gene can reveal its late functions if specialized, since organ identity will be specified by other players. As previously described, in *Phalaenopsis* flowers the lateral sepal identity is specified by the OAGL6-1/OAP3-1/OPI protein complex, but this organ also expresses OAGL6-2 very specifically in the lower part of the sepal, more prominently in a small region displaying red spots of pigmentation (Figure 1A). The knock-down of *OAGL6-2* by VIGS does not strongly affect lateral sepal identity since the OAGL6-1/OAP3-1/OPI complex is able to form, but strikingly, the red spots of pigmentation are lost (Hsu *et al*., 2021). This is clear evidence that *OAGL6-2* is necessary for the formation of these red spots, which only appear in a restricted domain of the sepal, likely because *OAGL6-2* expression is enriched there. A direct expectation from this result is that heterotopic expression of *OAGL6-2* in the whole lateral sepal should lead to the formation of red spots everywhere, but this remains to be tested. Similarly in Rice, the knock-out of *OsMADS2*, with distally enriched expression in the lodicule (Figure 1B), results in elongated lodicules, suggesting that *OsMADS2* acts locally to restrict their growth (Yadav *et al*., 2007; Zamzam *et al*., 2025). While the single knock-down of *OsMADS4* does not cause any lodicule phenotype, combining it with the *osmads2* mutation leads to severely flattened and elongated lodicules, increasing the effect of the single *osmads2* mutation. Therefore, and although *OsMADS4* is evenly expressed in the lodicule, it participates to the repression of lodicule elongation together with *OsMADS2*, itself with distally enriched expression. The target genes of OsMADS2 were recently identified by chromatin immunoprecipitation (ChIP-Seq) and RNA-Seq in the *osmads2* mutant, revealing potential key players in lodicule elongation, in particular related to water influx and accumulation of osmolites driving growth (Zamzam *et al*., 2025). A distal expression of these potential targets in the lodicule, and their knock-out phenotype, would support their role in its elongation. When examining pistil development, neither the single *osmads2* mutant nor the single *osmads4* knock-down line displayed any female fertility phenotype, but combining the two mutations affected embryo sac development and caused parthenocarpy (Zamzam *et al*., 2025). In that case, both genes appear fully redundant, although their expression pattern in the ovule is quite distinct (Figure 1B) and the role of the specific expression pattern of *OsMADS2*, if any, remains unclear.

Stage-specific knock-downs have the potential to elucidate the late and spatial-specific functions of floral MADS genes. For instance, Virus-Induced Gene Silencing (VIGS) experiments can be performed at late stages of flower development. Recently, VIGS against the B-class gene *PhDEF* (*Petunia x hybrida DEFICIENS*) on petunia flowers at anthesis revealed the late role of *PhDEF* in regulating scent production and emission, among others through the regulation of the key regulatory genes from the scent production pathway *EMISSION OF BENZENOIDS I* (*EOBI*) and *EOBII* (Bednarczyk *et al*., 2025). Interestingly, this regulation appears to be independent of PhDEF’s usual partners PhGLOBOSA1 (PhGLO1) and PhGLO2, that are essential for the transcriptional regulation activity of PhDEF at early stages of development, but might become accessory later (Manchado-Rojo *et al*., 2012; Bednarczyk *et al*., 2025). Transgenic approaches could also help explore the late role of floral MADS-box genes. For instance, pulsed knock-down of the B-class gene *APETALA3* (*AP3*) in Arabidopsis, by inducible expression of a microRNA targeting the gene, has confirmed the crucial early role of *AP3* in the specification of stamen identity, while petal identity is committed later (Wuest *et al*., 2012). Using a similar system at much later stages of development would allow testing the late role of AP3 in stamen and petal maturation. In Antirrhinum, the *def-101* mutant harbours a point mutation in the B-class *DEF* gene, and this mutation is temperature sensitive (likely due to an altered interaction of DEF with its partner GLO at high temperature (Zachgo *et al*., 1995). Shifting *def-101* plants between low and high temperatures allows to explore the early or late role of DEF in petal and stamen development (Zachgo *et al*., 1995). A late shift to high temperature leads to petals with green edges and stamens with ovules close to the base of stamen filaments, again confirming that B-class function is required until the very last stages of petal and stamen development for their correct morphogenesis (Zachgo *et al*., 1995). However, a detailed expression pattern of *DEF* is missing at this stage, that would link the localized effects of the mutation to the spatialized expression of *DEF*. Pulsed induction of expression at late stages of development is also possible with the GR-DEX inducible system, whereby application of dexamethasone results in the nuclear translocation of a MADS-GR fusion protein (Wellmer *et al*., 2006; Ó’Maoiléidigh *et al*., 2013). This approach revealed that prolonged activity of the C-class gene *AG* was necessary for stage-specific stamen organogenesis up until anthesis in Arabidopsis (Ito *et al*., 2007). One of the late functions of AG in stamens highlighted by this experiment was the late stage binding of AG to *DAD1 (DEFECTIVE IN ANTHER DEHISCENCE1)*, encoding a phospholipase A, catalyzing the first step of the jasmonic acid biosynthetic pathway necessary for stamen maturation (Ito *et al*., 2007). These types of assays, that could even be spatially restricted with the use of cell-type-specific promoters, and combined with a careful microscopic characterization of cell types and organ shape after late inactivation, constitute adequate tools to precisely evaluate the late role of floral MADS genes, in species amenable to genetic transformation.

Flower chimeras are another kind of genetic material that can reveal late and spatial-specific functions of floral MADS genes. Flower chimeras are composed of cells with more than one genotype (Frank and Chitwood, 2016). The chimeras discussed here, being a mixture of cells expressing and not expressing a floral MADS-box gene, can be either naturally observed as the result of transposition (*Tam3* transposon in Antirrhinum undergoing low-temperature-dependent transposition (Carpenter *et al*., 1987), or the highly active *dTph1* transposon in some cultivars of *Petunia x hybrida* (Gerats *et al*., 1990) or genetically engineered (using for instance the *Activator/Dissociator* transposable element system from maize, (Lazarow *et al*., 2013), or random X-ray mutagenesis (Bouhidel and Irish, 1996)). In the case of a transposable element that is inserted in a floral MADS gene, the flower will likely have a knock-out homeotic phenotype, but excisions of the transposon from the gene can result in restoration of gene function. Differences in the timing and position of the cell in which the excision happens, together with the rate and orientation of cell divisions within the organ, yields different kinds of chimeras (Frank and Chitwood, 2016). Periclinal chimeras will harbour different genotypes in their different cell layers (derived from the meristematic L1, L2 and L3 cell layers), while mericlinal chimeras will display sectors of wild-type cells within a mutant cell layer. Periclinal chimeras for the B-class homeotic genes *DEFICIENS* (*DEF*) and *GLOBOSA* (*GLO*) in Antirrhinum, *PhDEF* in petunia, and *AP3* and *PI* in Arabidopsis, have been characterized by ourselves and others (Bouhidel and Irish, 1996; Perbal *et al*., 1996; Efremova *et al*., 2001; Jenik and Irish, 2001; Vincent *et al*., 2003; Chopy *et al*., 2024). Overall, the phenotypes of these periclinal chimeras have revealed that B-class MADS-box genes can have different functions when active in one layer or the other of the petal. For instance, our own petunia periclinal *phdef* flowers strikingly show that *PhDEF* controls limb development and pigmentation from the epidermis, while it controls tube development from the mesophyll (Chopy *et al*., 2024). Later mericlinal chimeras, with small petal epidermal sectors expressing *PhDEF* among an otherwise *phdef* epidermal layer, further confirmed that late expression of *PhDEF* in the epidermis is sufficient to drive conical cell formation and pigmentation, while also driving limb growth locally, even at the anthesis stage (Figure 1H) (Chopy *et al*., 2024). We recently established that PhDEF indeed regulates a different set of target genes in the epidermis and in the mesophyll of the petal, with a major binding and regulatory action in the epidermis, even at a late developmental stage (Cavallini-Speisser, Désert *et al*., 2025, Preprint). If the flower chimeras can be induced, like in the case of the temperature-sensitive *Tam3* transposon, this has the potential to dissect the role of both time- and space-restricted expression of floral MADS genes. Flower chimeras can prove highly informative of late and spatialized roles of floral MADS genes, but this genetic material is difficult to obtain (either dependent on transposon insertion alleles randomly obtained, or to be genetically engineered, which is not accessible in non-transformable plant species) and to propagate, the chimeric phenotypes being non-heritable since gametes derive from the L2 layer exclusively.

In Box 1 and Figure 2, we detail the process of the megaspore mother cell (MMC) specification in the ovule of Arabidopsis flowers, since intricate mechanistic insights have recently been proposed to link the expression of the MADS-box genes *AG* and *STK* to their downstream players, in a precise spatio-temporal context. This example highlights the complexity of the regulatory processes involved in the specification of a precise cell type, given an initially broad expression pattern.

**Figure 2:**
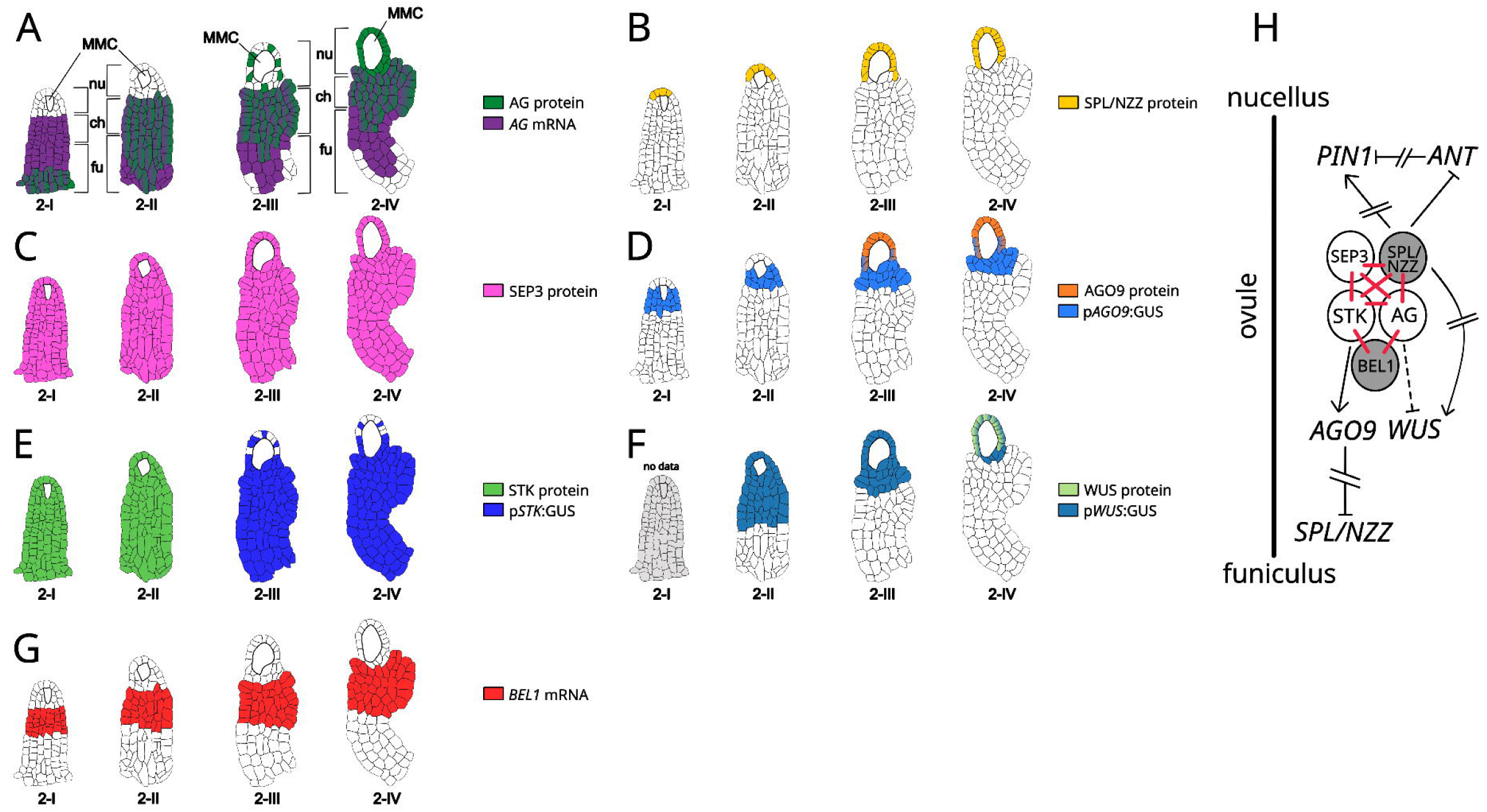
Localised expression of AG and STK in the ovule restricts downstream effectors for proper ovule specification. **(A)** *AG* transcript is detected in the funiculus and chalaza of the ovule from stages 2-I to 2-IV, slowly disappearing from the funiculus base. AG protein presence is gradually increasing in the nucellus from stages 2-III to 2-IV, suggesting a non-cell-autonomous action of AG in this cell layer. **(B)** The SPL/NZZ protein is restricted to the apical nucellar cells during stages 2-I and 2-II, then extends to the whole L1 nucellar layer in stages 2-III and 2-IV. **(C)** The SEP3 protein is expressed in all ovule tissues, except the MMC, from stages 2-I to 2-IV. **(D)** The *AGO9* transcript (as revealed by a GUS transcriptional reporter) is localized at the bottom part of the nucellus from stages 2-I to 2-IV, but the AGO9 protein is detected in the top nucellar layer as well in stages 2-III to 2-IV, and occasionally in the MMC (not shown). **(E)** In stages 2-I and 2-II, the STK protein is localized in all ovule tissues except the MMC, but the *STK* transcript still accumulates in the funiculus and chalaza, weakly in the nucellus, at later stages. The STK protein localization pattern was only available for stages 2-I/2-II. The *pSTK:GUS* expression pattern was only available for stages 2-III/2-IV. **(F)** From stages 2-II to 2-IV, *WUS* transcripts are gradually restricted from the chalaza and nucellus to the nucellus layer only, which is coherent with WUS protein being localized in the top nucellar layer at stage 2-IV. The *pWUS:GUS* expression pattern was not available for stage 2-I. The WUS protein localization pattern was only available for stage 2-IV. **(G)** *BEL1* transcripts are present only in the chalaza from stage 2-I to 2-IV. **(H)** Putative model of MADS-box proteins interactions in the ovule: SEP3, AG and STK can each individually interact with SPL/NZZ or with each other in the ovule, as well as with BEL1 for AG or STK dimers containing SEP3. STK binds and induces the expression of *AGO9*, that together with an indirect action through the RNA-directed DNA Methylation (RdDM) pathway, restricts *SPL/NZZ* expression to the top nucellar layer. SPL/NZZ in the apical nucellar layer represses *ANT* expression in the nucellus and indirectly induces *PIN1* expression, that establishes auxin fluxes important for MMC specification. In the same layer, *SPL/NZZ* promotes *WUS* expression that is also required for proper MMC specification. *WUS* expression is restricted to the nucellus probably by AG repressive action in the chalaza and the funiculus, in interaction with BEL1 that is specifically expressed in the chalaza. **Legends**. White circles represent MADS-box proteins, grey circles represent other proteins and red lines display the confirmed protein-protein interactions (by yeast two- or three-hybrid). Black arrows, black broken arrows, and black dotted arrows indicate a direct, indirect, or putative regulatory action, respectively. fu: funiculus; ch: chalaza; nu: nucellus; MMC: megaspore mother cell. Simplified ovule cellular organization derived from Vijayan *et al*., 2021. Ovule development stages after Schneitz *et al*., 1995. Transcript and protein expression patterns from (Reiser *et al*., 1995; Groß-Hardt *et al*., 2002; Urbanus *et al*., 2009; Olmedo-Monfil *et al*., 2010; Rodríguez-Leal *et al*., 2015; Zhao *et al*., 2017; Rodríguez-Cazorla *et al*., 2018; Cavalleri *et al*., 2025).

### Molecular mechanisms for spatial restriction of MADS-box gene function

#### Regulating MADS-box protein expression domains

In the previous paragraphs, we have reported multiple cases of MADS-box transcript or protein enrichment in specific parts of developing floral organs, sometimes in link with a localized function. The simplest mechanism to restrict floral MADS-box protein function to a specific location is indeed to restrict its presence there. The regulatory pathways that result in the establishment of the MADS-box gene expression domains in the floral meristem, evoked in the Introduction, are well established but are not necessarily the same ones that regulate the late expression pattern of these genes. For instance, activation of *AG* expression can happen independently of LFY at late stages of development (Weigel and Meyerowitz, 1993) and the auto-regulatory loop that is needed to maintain B-class gene expression is likely less important at late stages of development (Manchado-Rojo *et al*., 2012; Bednarczyk *et al*., 2025). Late maintenance of B- and C-class gene expression in Arabidopsis has been shown to rely on the action of the RNA helicase HUA ENHANCER2 (HEN2) (Western *et al*., 2002), together with the RNA-binding protein HUA1 and the putative transcription factor HUA2 (Chen and Meyerowitz, 1999; Li *et al*., 2001). The triple mutant *hua1 hua2 hen2* indeed shows correct expression of *AP3*, *PI* and *AG* in stamen and carpel primordia, but their expression becomes weak and patchy from stage 6 onwards, resulting in mosaic organs with altered identity. The identity of HEN2 as an RNA helicase suggests that this regulation happens post-transcriptionally, by controlling mRNA stability. Additionally, the putative RNA binding protein HUAENHANCER4 (HEN4), that interacts with HUA1 in the nucleus, acts specifically in the processing of *AG* pre-mRNA but not for other floral homeotic transcripts (Cheng *et al*., 2003). Post-transcriptional regulation of MADS-box genes is therefore a possible, but poorly described, mechanism to determine floral MADS genes expression patterns. Other pieces of evidence suggest that post-transcriptional regulation of MADS-box genes expression by small RNAs and long non-coding RNAs (lnc-RNAs) might be widespread: microRNAs miR444 and miR824 are involved in the regulation of MADS-box genes from the AGL17 family, with a role in regulating tillering and nitrate signalling in rice, and stomatal patterning and flowering time in Arabidopsis (Kutter *et al*., 2007; Gramzow *et al*., 2020). The lnc-RNA transcribed from *AG* second intron represses its expression in leaves, sepals and petals, while its absence leads to curly leaves and missing or misshaped sepals and petals, due to *AG* ectopic expression (Wu *et al*., 2018). In the diploid strawberry *Fragaria vesca*, four MADS-box genes are neighbours to lnc-RNAs, and their expression patterns correlate in specific tissues, suggesting a possible regulatory link, to be confirmed experimentally (Kang and Liu, 2015). A complex regulatory pattern involving transcription factors, chromatin remodelers, miRNAs, lnc-RNAs and RNA-binding proteins has been reported over the years for proper *AG* expression (Pelayo *et al*., 2021), suggesting that a similar picture might apply to other floral MADS-box genes, whose regulatory processes have been less studied.

Additionally to the regulation of MADS-box transcript levels or location, MADS-box protein location might also be regulated. For example, in spite of its homogeneous mRNA expression pattern in the petal, as previously described, the SEP3 protein is enriched in the adaxial epidermis of the petal side (Mandel and Yanofsky, 1998; Urbanus *et al*., 2009; Figure 1F). This suggests that either *SEP3* mRNA translation is higher in the adaxial epidermis than in the other cells of the petal, or that the SEP3 protein is degraded in the abaxial epidermis, or that the SEP3 protein moves between cells to accumulate in the adaxial epidermis. One possible way to discriminate between those two mechanisms is to prevent MADS-box protein movement, for instance by fusing it to a large protein (typically 3 GFP in tandem) that renders it unable to move through plasmodesmata, which should affect its distribution only if cell-to-cell movement is the main driving mechanism. To our knowledge, these experiments have not been performed with floral MADS-box proteins. Alternatively, the use of proteasome inhibitors should inhibit protein degradation but not protein movement; in the case of SEP3, such inhibitors did not alter the pattern of accumulation of SEP3, suggesting that protein degradation is not a major player in this process (Urbanus, 2010). Overall, it is generally assumed that the discrepancy between mRNA and protein expression patterns is due to protein movement. SEP3 is not the only protein suspected of moving between cells: the *AG* transcript is detected in the ovule integuments and not in the nucellus, but the *pAG::gAG*:*GFP* fusion is expressed in the nucellus, suggesting that the AG protein could move from the integuments to the nucellus (Reiser *et al*., 1995; Urbanus *et al*., 2009, and note the striking discrepancy between *AG* transcript and AG protein expression patterns in the pistil, Figures 1F, G). Consistently, AG was shown to move across cell layers when expressed only in the epidermis (*pL1::gAG*:*GFP*) in the floral meristem, leading to the suppression of *WUS* expression in the center of the floral meristem (Urbanus *et al*., 2010). In the previously described periclinal chimeras of DEG and GLO in Antirrhinum flowers, immunolocalization revealed that the DEF and GLO proteins expressed in the L2 and L3 layers were able to move to the epidermis in small amounts, while the converse movement was not observed (Perbal *et al*., 1996). MADS-box proteins thus appear to be able to move between cells, but this seems to be restricted to specific contexts and the full extent of this phenomenon remains to be characterized. Additionally, regulation of the cytoplasmic or nuclear localization of MADS-box proteins might act as another mechanism to regulate their function. Indeed, it was clearly observed that SEP3 subcellular localization changes very dynamically over the course of floral organ development, with for instance an initial nuclear localization in the inflorescence and flower meristem, and a cytoplasmic and nuclear localization in future petals and stamens (Urbanus *et al*., 2009). It was also observed that AP3 and PI only translocate to the nucleus when they dimerize (McGonigle *et al*., 1996), and a similar behaviour was reported for other MADS-box proteins (Immink *et al*., 2002; Bemer *et al*., 2008). Cytoplasmic accumulation might be a mechanism by which cells have a reservoir of MADS-box proteins, quickly available by nuclear translocation to exert their regulatory action for proper floral organ development; it might be linked with intercellular transport since proteins have to be in the cytoplasm to move between cells; and it might also be a transient accumulation of proteins before they are degraded (Urbanus *et al*., 2009).

#### Achieving spatialized functions with non-spatialized expression

Even with homogeneous protein expression within a floral organ, it is still possible that a floral MADS-box protein could play different functions in different regions of this organ. Cells being in different chromatin contexts, access to individual binding sites might be different between *i.e.* the distal and proximal region of an organ. These cell-specific chromatin profiles can be revealed by single-cell studies, such as single-cell ATAC-Seq (Assay for Transposase-Accessible Chromatin). A recent study on rice and sorghum leaves evidenced distinct patterns of chromatin accessibility in the epidermis and mesophyll (Swift *et al*., 2024). Since floral organs are modified leaves, and epidermal identity is established early during embryo development (Robinson and Roeder, 2015; Schrick *et al*., 2023), it is reasonable to assume that differential chromatin accessibility between the epidermis and the mesophyll is also true in floral organs. Additionally, MADS-box proteins themselves interact with, and regulate the expression of, chromatin remodelers (Smaczniak *et al*., 2012; Abraham-Juárez *et al*., 2020; Pelayo *et al*., 2021; van Mourik *et al*., 2023), suggesting that they could also modify pre-existing chromatin profiles and reinforce the specification of cell types.

Floral MADS-box proteins might also have access to different protein partners in different cell contexts. The famous floral quartet model proposed that different MADS-box protein complexes bind to different binding sites and regulate the expression of different genes (Theißen and Saedler, 2001). This applies to floral organ identity determination, but also to developing floral organs, in which different MADS-box protein complexes might form at different locations and fulfill different functions. Again, the case of orchid petals supports this mechanism, since the L’ complex (OAGL6-2/OAP3-1/OPI) forms at the lower and proximal part of the lateral sepal, resulting in the formation of red spots, while the SP complex (OAGL6-1/OAP3-1/OPI) forms in the rest of the lateral sepal (Hsu *et al*., 2021, Figure 1A). The identity of MADS-box protein partners has been investigated at a large scale by co-immunoprecipitation coupled to mass spectrometry, for instance in Arabidopsis flower buds and pistils, and in maize tassels (Smaczniak *et al*., 2012; Abraham-Juárez *et al*., 2020; van Mourik *et al*., 2023). These partners extend well beyond other MADS-box proteins and encompass a vast diversity of transcription factors, chromatin remodelers, enzymes and proteins of various functions, opening the possibility of many different complexes being formed in different cellular contexts. FUL has a dual role in inflorescence meristem specification and pistil/fruit development, that are temporally uncoupled (Ferrándiz *et al*., 2000b,*a*; van Mourik *et al*., 2023). This dual role is subtended by the formation of different protein complexes, in particular different FUL-MADS dimers, that bind to distinct binding sites and regulate the expression of different target genes, with some overlap between dimers (van Mourik *et al*., 2023). The different DNA binding specificities of FUL dimers alone are enough to predict most of their specificity of action, suggesting that other aspects, like chromatin accessibility, are likely minor players for the dual role of FUL (van Mourik *et al*., 2023).

### Conclusion

Floral homeotic MADS-box genes, after specifying early floral organ identity, can display highly complex and dynamical expression patterns, specific to some regions or cell types of floral organs. We have attempted to illustrate the diversity of these patterns, and while most patterns are diverse between species and classes of genes, we have also noted that B-class genes systematically display distally-enriched expression in the petal, often associated with epidermal enrichment. Although the characterization of the late functions of MADS-box genes is often inaccessible due to their early role in specifying floral organ identity, some studies were able to demonstrate the importance of their late expression for specific cell types. The potential molecular mechanisms governing this spatial restriction of MADS-box gene function are multiple and non-exclusive: epigenetic, transcriptional and post-transcriptional control of MADS-box gene expression and protein localization, differential chromatin status at target genes in different cell types, tissue-specific proteins partners, etc. How these different molecular mechanisms are deployed in specific cell types remains to be explored at a large scale. The advent of single-cell technologies, such as single-cell ChIP-Seq for histone marks, single-cell DNA methylation profiling, single-cell proteomics and of course single-cell or -nuclei RNA-Seq, have the potential to explore the cell-specific contexts that floral MADS-box proteins might encounter in a given cell type, and how it might influence their function.

## Author Contributions

E.D. and M.M. conceptualized, wrote, reviewed and edited the manuscript. M.V. reviewed the final version of the manuscript.

## Acknowledgements

We thank Richard Immink for discussions on MADS proteins movement.

## Generative AI statement

No generative AI was used in the making of this manuscript.

## BOX 1: A case study: from AG and STK localized expression to proper ovule specification

In Arabidopsis, ovules originate from the placental tissue and form protrusions during floral developmental stages 8 and 9. Because ovule development is not synchronous with floral development after stage 12, ovule-specific developmental stages have been defined (Schneitz *et al*., 1995). The expression patterns of MADS-box genes implicated in ovule development and other genes of interest span most of stage 2 of ovule development (2-I to 2-IV), after primordia initiation and elongation in stage 1-I and 1-II. At stage 2-I, the ovule is composed of three different domains: the funiculus, the chalaza and the nucellus containing one megaspore mother cell (MMC), which will enlarge through stages 2-I to 2-IV, while inner and outer integuments will initiate from the chalazal part of the ovule starting stages 2-II and 2-III respectively (Schneitz *et al*., 1995). MMC meiosis results in the production of a tetrad of haploid megaspores at stage 2-V, among which only one will differentiate into the female gametophyte: the embryo sac (Schneitz *et al*., 1995; Pinto *et al*., 2019).

Several MADS-box genes redundantly specify ovule identity in Arabidopsis: *AG*, *SEEDSTICK* (*STK*), *SHATTERPROOF1* (*SHP1*) and *SHP2*, together with the *SEP* genes (Favaro *et al*., 2003; Pinyopich *et al*., 2003). *AG* and STK are first expressed in the funiculus, chalaza and future integuments, and their expression gradually increases in the nucellus (Figure 2A and E; Urbanus *et al*., 2009; Rodríguez-Cazorla *et al*., 2018; Cavalleri *et al*., 2025), while *SEP3* is homogeneously expressed in the ovule throughout stages 2-I to 2-IV (Figure 2C; Urbanus *et al*., 2009). The downstream players of interest for MMC specification are WUS, that together with its downstream effectors peptides like WINDHOSE1 and WINDHOSE2, act to promote MMC formation (Figure 2G; Lieber *et al*., 2011; Pinto *et al*., 2019), and SPOROCYTELESS/NOZZLE (SPL/NZZ), a SPEAR-family transcription factor crucial for MMC differentiation capable of dimerizing and interacting with SEP3, AG, SHP1/2 and STK, as evidenced by yeast two- or three-hybrid assays (Schiefthaler *et al*., 1999; Yang *et al*., 1999; Brambilla *et al*., 2007; Cavalleri *et al*., 2025).

The expression of the WUS and SPL/NZZ proteins is restricted to the apical nucellar layer, very early for SPL/NZZ and later for WUS (Figure 2B and F). For WUS, this restricted expression pattern could be due to the joint action of AG, that is a well-established repressor of *WUS* transcription (Lenhard *et al*., 2001; Lohmann *et al*., 2001; Liu *et al*., 2011), and of the homeodomain transcription factor BEL1 expressed only in the chalaza (Figure 2G,Brambilla *et al*., 2007). Indeed, in *bel1* ovules, *WUS* expression is extended from the nucellus into the chalaza and funiculus (Brambilla *et al*., 2007). BEL1 was shown to interact with MADS-box dimers composed of SEP1 (or SEP3) and either AG, SHP1 or SH2, but only the interaction with AG seems to be important for *WUS* repression in the chalaza as *WUS* expression pattern is normal in the *stk shp1 shp2* triple mutant (Brambilla *et al*., 2007).

For *SPL/NZZ*, the restriction of its expression domain to the L1 nucellar layer is due to the indirect action of STK, potentially through regulation of the RNA-directed DNA Methylation pathway (RdDM) in the basal nucellar cells. Actors of the RdDM pathway include ARGONAUTE proteins (AGO), like AGO9, that can use small interfering RNAs (siRNAs) to silence transposable elements or genes. In *stk* and *ago9* ovules, additional MMCs are formed due to the ectopic expression of *SPL/NZZ* in the whole nucellus (Mendes *et al*., 2020). *AGO9* expression is downregulated in *stk* ovules, and it was shown by ChIP that STK can directly bind to *AGO9* and drive its expression in the ovule (Mendes *et al*., 2020). *STK* transcriptional activity is more pronounced in the basal nucellar cells than in the apical nucellar cells at stages 2-III and 2-IV of ovule development, in correlation with *AGO9* transcriptional activity in the same domain (Figure 2D,E). The AGO9 protein also partially colocalizes with SPL/NZZ in the L1 nucellar cells (Figure 2B,D), supporting the role of STK in restricting SPL/NZZ localization (Figure 2G). *WUS* expression maintenance in the nucellus is also thought to depend on SPL/NZZ positive regulation (Sieber *et al*., 2004). In summary, the combined action of AG, STK, BEL1 and AGO9, together with secondary players like SEP3 and SHP1/2, are crucial to restrict WUS and SPL/NZZ to their correct expression domains. Failure to do so results in the expression of chalazal/integument identity transcription factors like *AINTEGUMENTA (ANT)* in the nucellus, and improper control of auxin fluxes through misregulation of the auxin transporter-encoding gene *PIN1*, resulting in defective MMC formation by non-cell-autonomous mechanisms (Cavalleri *et al*., 2025). Altogether, this example (that we have simplified for the purpose of this review) illustrates the complex pathways that lead from broad MADS-box gene expression to restricted expression of downstream effectors for a cell-type-specific function.

